# The mini yet mighty stapes: a comparison of ancient DNA yields among ossicles and the petrous bone

**DOI:** 10.1101/2025.07.17.664655

**Authors:** Ekin Sağlıcan, Arda Sevkar, Duygu Deniz Kazancı, Sevgi Yorulmaz, Kıvılcım Başak Vural, Gökhan Çakan, Gökçen Adiloğlu, Güneş Duru, Brenna Hassett, Nurcan Kayacan, Aslı Erim Özdoğan, Anders Götherström, Füsun Özer, Ömür Dilek Erdal, Yılmaz Selim Erdal, Mehmet Somel

## Abstract

The petrous bone is considered the most efficient source of endogenous DNA across skeletal tissues in ancient DNA research as well as in forensic work. Recently, ancient DNA (aDNA) in auditory ossicle bones was shown to be comparably well-preserved as in the petrous, although no attempt was made to distinguish among the three ossicle bones. In this study, we used a total of 114 human ossicle- and petrous-derived sequencing libraries from similar contexts (c.10,000 BP - 7,000 BP Anatolia), including 34 matched libraries prepared from the same individuals’ ossicle and petrous bones. Our results suggest that endogenous human aDNA preservation in the stapes is on average two times higher than in the petrous bone; it also tends to be higher than in the malleus and incus. Similarly, aDNA fragment lengths were higher in the stapes than in the petrous, whereas postmortem damage, clonality and contamination rates were comparable. Despite being the smallest bone in the human skeleton, the stapes may be the most optimal aDNA source yet identified.

## Introduction

The efficient retrieval of postmortem or ancient DNA (aDNA) from skeletal remains is a major limiting factor in paleogenomics and forensic genetics. Over the last decade, the petrous bone has been recognised as the most effective source in both archaeological and forensic settings (Gamba et al., 2014; Gaudio et al., 2019; Pinhasi et al., 2015, 2019). Endogenous DNA preservation in the petrous bone (and more specifically, the osseous labyrinth, or the otic capsule, which includes the cochlea) tends to be higher than in other skeletal elements, including long bones (femur, tibia etc.), phalanges, teeth, calcaneus, and the talus (Haarkötter et al., 2023; Hofreiter et al., 2021; Parker et al., 2020; Zupanič Pajnič et al., 2025). This property is attributed to limited bone remodelling and a high concentration of osteocytes in the petrous (İbrahim et al., 2022; Pinhasi et al., 2015).

Before the discovery of the petrous bone as an effective aDNA source, Bell and colleagues had suggested that ear ossicles, which contain a high proportion of mineralised osteocytes, could be an optimal target for DNA and protein retrieval from postmortem material (Bell et al., 2008). More recently, Sirak *et. al*. (2020) compared ancient DNA yields in ossicles and petrous bone from 10 human skeletons by shotgun sequencing. These authors found: a) similar endogenous DNA content in ossicles as in the petrous bone (with non-significantly higher levels in ossicles), b) slightly lower DNA postmortem damage (in the form of cytosine deamination) in ossicles, c) three times higher human contamination in ossicles (even though all the libraries studied had <5% contamination levels), and d) similar levels of library complexity.

Sirak and colleagues thus concluded that ossicle bones could be a useful source of aDNA, especially when the petrous bone is not accessible and/or when its destructive sampling is undesirable, as its morphology can be used for studying genetic relationships (Sirak et al., 2020). A few paleogenomic and forensic genetic studies have also included ossicles as a DNA source, although with limited sample sizes and/or without comparison with the petrous bone (Prendergast et al., 2019; Schwark et al., 2015). Meanwhile, no study, to the best of our knowledge, attempted to differentiate between the three different auditory ossicle elements: the stapes, malleus, and incus. These three bones might be expected to differ in DNA preservation levels given the differences in their developmental profiles, bone and osteocyte densities (Ivanovic et al., 2024; Marotti et al., 1998; Richard et al., 2017).

## Results

Here we compared aDNA sequencing libraries obtained from 17 archaeological skeletons with both petrous and ossicle tissue, including 7 stapes, 5 malleus, 2 incus, and 3 unidentifiable ossicle fragments **(Figure 1A; Supplementary Figure 1)**. The skeletons included adults and subadults, and derive from 4 archaeological excavation sites from across ancient Central Anatolia and Upper Mesopotamia, ranging from c.10,000 BP to 7,000 BP, and covering mainly temperate steppe-like environments (**Supplementary Table 1**; **Supplementary Figure 2**). From the site of Köşkhöyük, where the majority of our samples derive from, we additionally expanded our dataset with aDNA data from 5 stapes, 9 malleus, 4 incus and 4 unidentifiable ossicle fragments that did not have matching petrous libraries; we also used 69 petrous-derived libraries that did not have matching ossicle libraries **(Supplementary Table 1)**.

**Figure 1:**
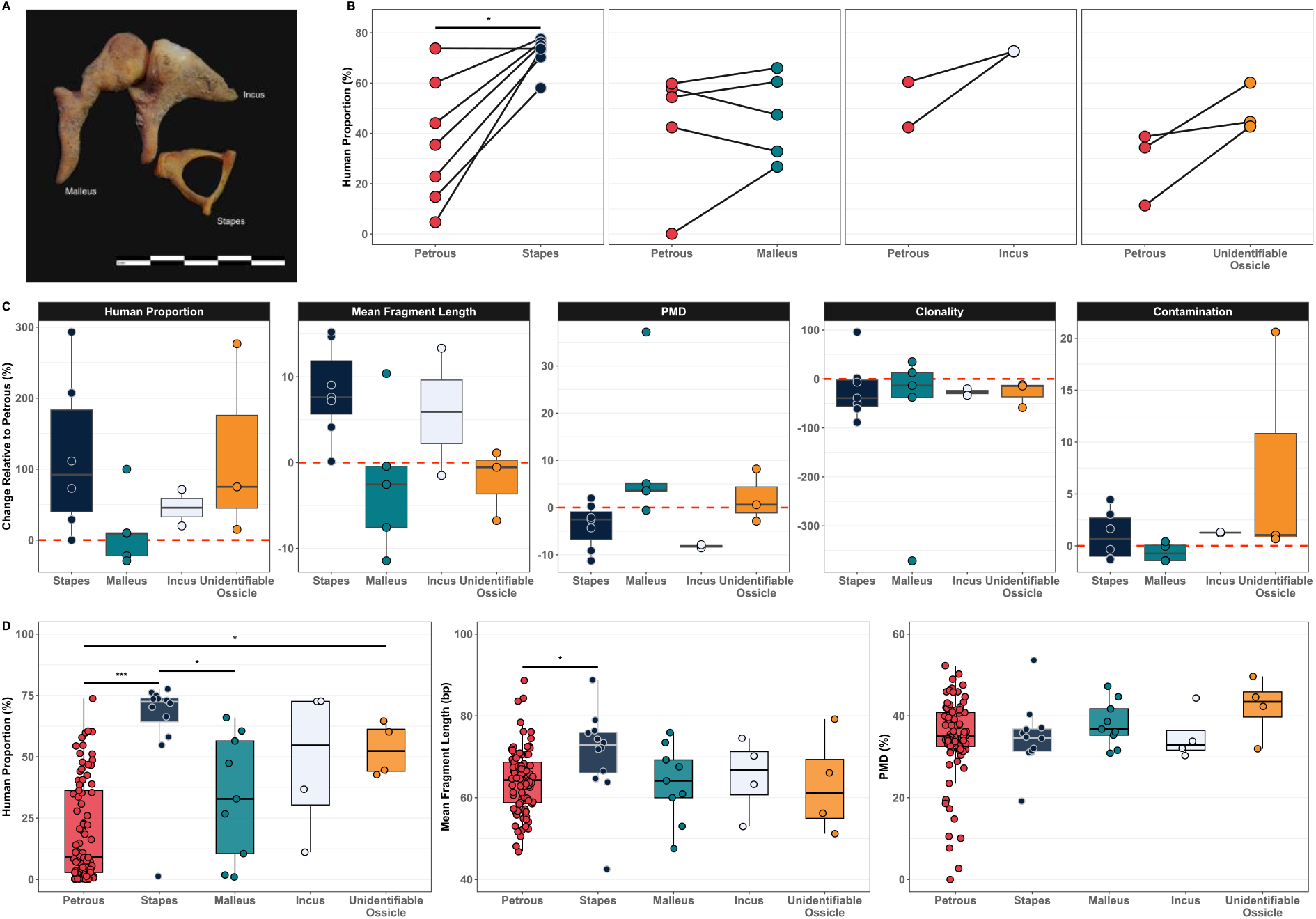
Auditory ossicle bones and their comparison with the petrous bone as a source of ancient DNA. **A)** Representative auditory ossicle bones used in this study. Each black and white segment in the scale bar represents 2 millimeters. **B)** Comparison of the human DNA proportion in shotgun DNA libraries prepared from each ossicle type relative to those prepared from the petrous bone. Libraries derived from the same individual are connected by lines. **C)** Change in sequencing metrics for ossicle bones relative to petrous bones. One outlier (individual bah037, ∼1400% increase) was excluded from the plot for visualization clarity. PMD: postmortem damage. **D)** Sequencing metrics for all libraries, including those with and without matching petrous or non-petrous counterparts. Only statistically significant Wilcoxon signed-rank (WSR) test results are shown above boxplots (*: p<0.05; ***: p<0.001).

The ossicles were dissolved directly in the extraction buffer, while the petrous bone samples were pulverised by drilling, and DNA was extracted using the Dabney protocol (2013) **(Methods)**. We constructed dual-indexed, double-stranded Illumina libraries (Meyer & Kircher, 2010) and shotgun sequenced each library once between approximately 4 - 60 million reads (median ∼12.5 million) **(Supplementary Table 1)**. We mapped the data to the human reference genome (hg19) and compared various characteristics of the ossicle- and petrous bone-based libraries **(Methods)**: human proportion, fragment length, postmortem damage (PMD), clonality, and contamination.

Endogenous human DNA proportion, an indicator of DNA preservation, was calculated as the fraction of reads mapping to the human genome (without removing duplicates, to avoid confounding by sequencing depth differences among libraries). We found a clear enrichment for the stapes relative to the petrous bone in endogenous human DNA **(Figure 1B-C, Supplementary Figure3A)**. In 6 of 7 comparisons, the stapes had higher values with a median gain of ∼111%. This was overall a significant increase [Wilcoxon signed rank (WSR) test p=0.031]. The only case where we saw practically no difference between the petrous and the stapes already had ∼70% human endogenous DNA in both tissue samples.

Among the incus, malleus, and unidentified ossicle fragments, 8 in 10 comparisons revealed higher endogenous human DNA than the petrous, although the gains were more modest compared to that observed with the stapes (median ∼11%; p=0.01) (**Figure 1B-C, Supplementary Figure3A**). Overall, across the combined set of 17 ossicle-based versus petrous-based libraries, there was a visible and significant increase in endogenous human DNA in the former (p=0.008).

We further compared DNA content across all 114 ossicle and petrous-derived libraries in our expanded dataset (stapes: 12, malleus: 9, incus: 4, unidentifiable ossicle: 4 and petrous: 85, belonging to either the same or to different individuals). Of these, 104 were from the Neolithic/Chalcolithic levels of the site of Köşkhöyük. These comparisons also supported significantly higher DNA preservation in the stapes than all other studied tissues (Mann-Whitney U (MWU) test p<0.05), except for the incus for which our sample size was too low (**Figure 1D**).

Human DNA fragment length is another indicator of DNA preservation. Fragment lengths in the 7 stapes-derived libraries were on average significantly longer (mean 5.5 bp) than matching petrous-derived ones (WSR test p=0.02; **Figure 1C, Supplementary Figure3B**). The same pattern was observed across the set of all 114 ossicle and petrous libraries: we found longer fragments in the stapes compared to the petrous (**Figure 1D**, MWU test p=0.05). The other ossicle tissues did not differ from the petrous in fragment length (**Figure 1C-D**, p>0.05).

We next studied PMD, estimated as cytosine deamination-caused C-to-T mismatches on fragment ends. We found similar PMD levels between ossicles and matched petrous libraries for each ossicle type (WSR test p>0.05; **Figure 1C, Supplementary Figure3C**).

Sirak and colleagues had reported slightly but non-significantly higher clonality (inverse of library complexity) and also significantly higher contamination levels in their ossicle libraries compared to their petrous libraries (Sirak et al., 2020). We measured clonality as the proportion of duplicated reads in libraries randomly downsampled to 1 million reads each. Clonalities of ossicle-derived libraries were similar to those of matched petrous ones, with no significant difference (**Figure 1C, Supplementary Figure 3D**, WSR p>0.05). Further, using the *contamMix* method (Fu et al., 2013), we did not find any significant difference in human contamination levels between ossicle- and petrous-based libraries (**Figure 1C**, WSR p>0.05).

## Discussion

Sirak *et al*. had previously suggested that ossicles could be used as an effective alternative to the petrous bone as an ancient DNA source (Sirak et al., 2020). Our results now indicate the stapes as a significantly better source of aDNA than the petrous, and possibly also than other ossicles. This quality is reflected not only in higher endogenous DNA but also in the longer DNA fragments found in stapes-based libraries. This would potentially elevate the stapes to the top of the list of human skeletal elements for ancient and forensic DNA retrieval.

Our conclusions are subject to a number of limitations. First, the incus was represented by only four samples in our study, so we could not fully explore its potential as aDNA source. Second, we extracted DNA from the petrous bone using drilling instead of cutting and dissolving; the latter was the protocol Sirak *et al*. (2020) performed and may be more efficient in retrieving petrous DNA than our approach. However, like Sirak *et al*. (2020), we find similar endogenous DNA proportions between the petrous and non-stapes ossicles. Our ossicle processing approaches are also the same as that study. Together, these suggest that the higher performance of the stapes relative to both the petrous bone and to other ossicles cannot be fully explained by putative low DNA retrieval efficiency in our petrous libraries caused by drilling.

Another possible explanation for the observed differences between our study and Sirak *et al*. (2020) may be the age of the individuals included. Sirak *et al*. did not report the ages of their sample, but all except one of the individuals with stapes samples studied here were subadults (mainly neonates and infants, due to the nearly exclusive representation of subadult burials at Köşkhöyük). It is plausible that a rapid reduction of osteocytes in the stapes by apoptosis that occurs within the first year of life and continues at a lower pace at later ages (Rolvien et al., 2018) could lower retrievable DNA levels in the stapes of adults. Countering this, we note that the only adult individual (bah037) in our dataset with both stapes- and petrous-based libraries had the strongest DNA preservation difference, implying that age may not be a relevant factor. Finally, all our comparisons derive from archaeological sites from early Holocene Southwest Asia, although Sirak *et al*. used a broader geographic range. Nevertheless, it appears unlikely that the high performance of the stapes would be particular to samples from a specific period and region.

The ossicle bones are known to undergo only limited remodelling after 1-2 years of age (Marotti et al., 1998; Rolvien et al., 2018), a pattern also observed in the petrous bone (Frisch et al., 1998; Sølvsten Sørensen et al., 1992). Sirak and colleagues (2020) argued that this limited remodelling could explain comparable DNA preservation levels in ossicle and petrous samples. In the absence of remodelling, mineralised, apoptotic-like osteocytes with condensed chromatin structures are reported to accumulate in ossicles (Bell et al., 2008; Rolvien et al., 2018), which could be the source of preserved aDNA. The reason for the relatively high aDNA preservation in the stapes we observe here is yet unclear, but various studies have noted differences in histology and development among ossicles, including the later ossification of the stapes compared to the malleus and incus (Richard et al., 2017), high bone density in the stapes footplate (Ivanovic et al., 2024), and a higher percentage of dead osteocytes in the stapes compared to the other ossicles through lifetime (Marotti et al., 1998).

Our work thus indicates that the stapes, with its small size (c. 3-6 mm) and fragile appearance, can be a superior source of DNA over the petrous as well as any other human bone element, at least in the skeletons of young individuals, but likely also in those of adults. The stapes could therefore become the target tissue for archaeogenomic studies and also in forensic analysis. The stapes also has the advantage of being readily retrievable without destruction to the temporal bone. We thus echo the call by Sirak and colleagues to the archaeology and bioarchaeology communities to treat skull fragments carefully to avoid the loss of ossicles during excavation and further processing.

## Methods

All experiments were conducted in dedicated ancient DNA laboratories at Middle East Technical University and Hacettepe University in Ankara, Türkiye. Several precautionary measures were taken to minimise the risk of contamination during the experiments. All equipment, utensils and laboratory surfaces were decontaminated with DNA AWAY (Thermo Fisher) or a bleach solution before each use and in-between different samples. All solutions not sensitive to UV light were exposed to UV-irradiation for 30 minutes in a UV crosslinker. Negative controls were included at every step necessary to monitor potential contamination.

### Sampling and aDNA extraction

We used a modified protocol based on Dabney et. al. (2013) for ancient DNA extraction. The auditory ossicle bones were directly added into the extraction buffer, without pulverising or cleaning. From the petrous bone (otic capsule) material, we obtained approximately 80 mg of bone powder by drilling into surface-cleaned (using 1% bleach solution followed by rinsing with sterile water) bones. All samples (ossicles or petrous bone powder) were incubated in 1 ml of extraction buffer (0.45M EDTA and 0.25 mg/ml Proteinase K, pH 8) at 37°C in a rotor incubator. After 24 hr incubation, we centrifuged the tubes and transferred the supernatants into new clean tubes without touching the pellet. Then, an additional 1 ml of extraction buffer was added onto the pellet and incubated again at 37°C for 24 hrs. After centrifuging the tubes, the supernatants from the two digestions for each sample were combined. Next, we added 13 ml of freshly prepared binding buffer (pH 5.2) (5M Guanidine Hydrochloride, 40% (vol/vol) Isopropanol, 0.05% Tween-20 and 90 mM Sodium Acetate) onto the supernatants and filtered the mix through MinElute Silica Columns (Qiagen). Silica columns were washed twice with 750 μl PE buffer (Qiagen). At the final step, 50 μl of EB buffer (Qiagen) was added onto the filters in silica columns for DNA elution.

### Library preparation and indexing

We followed Meyer and Kircher (2010) protocol for preparing Illumina-compatible double-stranded libraries. The procedure has three main steps: blunt-end repair, adapter ligation, and adapter fill-in, with two MinElute silica column (Qiagen) purification steps in between. First, the blunt-end repair of aDNA fragments was performed with T4 DNA polymerase (Thermo Fisher Scientific), followed by the first MinElute silica column purification step. Then, using T4 DNA Ligase (Thermo Fisher Scientific), adapter oligos were ligated to the blunt-end repaired aDNA fragments from the previous step. Adapter-ligated DNA molecules were purified using MinElute silica columns. Finally, the adapter fill-in step was performed with *Bst* Polymerase, Large Fragment (New England Biolabs). To check the quality of the double-stranded libraries, we performed qPCR analysis. The qPCR Ct values were used to estimate the number of PCR cycles necessary to amplify each library to the threshold value. Lastly, each library was amplified and barcoded with Illumina-compatible indexes in three 50 μl PCR reactions (Kircher et al., 2012).

### Size selection of aDNA fragments and purification of the libraries

For size selection and purification, three PCR reactions (∼150 µl total) per library were pooled in low-binding microcentrifuge tubes. AMPure XP beads (75 µl) (Beckman Coulter), equilibrated to room temperature, were added to PCR products at a 0.5X ratio (AMPure XP beads:PCR product) to selectively bind larger DNA fragments. The mixture was vortexed, briefly pulse-spun, and incubated at room temperature for 5 minutes before being placed on a magnetic rack until the beads aggregated (∼1 minute). The supernatant, containing smaller fragments, was carefully transferred to a new tube. A second round of size selection was performed by adding AMPure XP beads at a 1.8X ratio (270 µl) to the recovered supernatant, followed by vortexing, brief pulse-spinning, and incubation for 10 minutes at room temperature. After placing on the magnetic rack (∼3 minutes), the supernatant was discarded and the beads bound to the target DNA, were washed three times with 200 µl of freshly prepared 70% ethanol. Each wash included vortexing, pulse-spinning, and magnetic separation. Following the final wash, beads were air-dried until residual ethanol evaporated. DNA was then eluted in 36 µl of TET buffer (TE + 0.05% Tween-20), vortexed for 20 seconds, pulse-spun, and incubated for 5 minutes at room temperature. A final magnetic separation for 5 minutes allowed recovery of the eluate, which was transferred to a clean low-binding tube for downstream analyses.

### Quality control of libraries and sequencing

The concentrations of purified libraries were measured with the Qubit dsDNA HS Kit. Highly concentrated libraries were diluted to ∼2 ng/μl. Next, the fragment length and concentration of the libraries were assessed using an Agilent 2100 Bioanalyzer DNA High Sensitivity Kit. Libraries with fragment lengths between 150 and 300 bps passed the quality check. For sequencing, libraries with unique index pairs were pooled in equimolar concentrations (final concentration of 10 nM total pool) and sequenced 2×150 cycles on Illumina Novaseq 6000 S1/S4 platforms. Each library was sequenced once.

### Bioinformatic analyses

Adapter sequences were removed using “*-qualitybase 33 -gzip -trimns*”’ parameters by using the *AdapterRemoval* tool (Schubert et al., 2016). The libraries, sequenced by paired-end reads, were merged after removing residual adapter sequences, requiring at least 11 bp overlap between the pairs using additionally “*-collapse -minalignmentlength 11*”. The quality of the sequences were studied using the *FASTQC* software (*https://www.bioinformatics.babraham.ac.uk/projects/fastqc/).* We obtained the sequencing number from these FASTQ files. Libraries were aligned to the human reference genome (*v*.*hs37d5*) using the “*aln/samse*” algorithm of the *Burrows-Wheeler Alignment (BWA)* software (*v 0*.*7*.*15*) (Li et al., 2009) with parameters “*-n 0*.*01, -o 2*” and by disabling the seed with “*-l 16500*”. We calculated the human proportion using the BAM files obtained as a result of this alignment process. PCR duplicates with identical start and end coordinates were removed using the script “*FilterUniqueSAMCons*.*py*” (Kircher, 2012). To minimise spurious alignment, reads with alignment quality of less than 30, shorter than 35 base pairs, and containing more than 10% mismatch were discarded.

After alignment, we calculated the average genome coverage using the *genomeCoverageBed* algorithm implemented in the *bedtools2* software (Quinlan & Hall, 2010) with reads with a mapping quality >30. The aligned libraries were examined using the *PMDtools* algorithm with the “*-deamination*” parameter to calculate postmortem damage in DNA (Skoglund et al., 2014). The base position was determined where the average postmortem damage (cytosine to thymine deamination) at the 5’ ends of reads in each library falls below 1%. The distribution of read lengths and contamination probabilities based on mitochondrial DNA diversity were calculated with the *contamMix* algorithm using default parameters (Fu et al., 2013).

To determine the human endogenous DNA content of each obtained library, we used the ratio of the sequences (i.e. merged reads) aligned to the human reference genome to all sequences in that library. We used the filtered sequences (as described above) but without removing duplicates (because technical factors such as higher PCR cycles or sequencing depth will increase duplication levels).

To estimate clonality, we downsampled each BAM file to 1 million reads. Downsampling was then performed using the “*-s*” parameter in samtools (Li et al., 2009) with the calculated fraction. The percentage of clonality, which we define as the proportion of duplicate reads to total reads, was calculated for each downsampled BAM file using the same procedure described above.

## Statistical analyses

All statistical analyses were conducted in R (R Core Team, 2025). We used the packages *readxl* (Wickham & Bryan, 2023), tidyverse (Wickham et al., 2019), and *reshape2* (Wickham, 2007) for data processing. Mann-Whitney U and Wilcoxon signed-rank tests were performed using the *wilcox*.*test* function from the R *stats* package (R Core Team, 2025). Visualizations were created using *ggplot2* (Villanueva & Chen, 2019), *superb* (Cousineau et al., 2021), *rnaturalearth* (Massicotte & South, 2023), *raster* (Hijmans, 2023), *ggrepel* (Slowikowski, 2024), *ggpubr* (Kassambara, 2023), *ggforce* (Pedersen, 2024), *png* (Urbanek, 2022), *grid* (R Core Team, 2025), *ggh4x* (Brand, 2024), *DescTools* (Signorell, 2024) R packages.

## Code Availability

All R codes related to the statistical analyses of this project are available in the following GitHub repository: https://github.com/ardasevkar/Auditory-Ossicles.

## Acknowledgements

We thank members of the CompEvo (Middle East Technical University) and HUMANG (Hacettepe) groups, Eva-Maria Geigl, Thierry Grange, Vendela Kempe Lagerholm, N. Ezgi Altinişik, Hannah Moots and Aliye Öztan for support and helpful suggestions.

