## Supplementary Figures for "The mini yet mighty stapes: a comparison of ancient DNA yields among ossicles and the petrous bone"

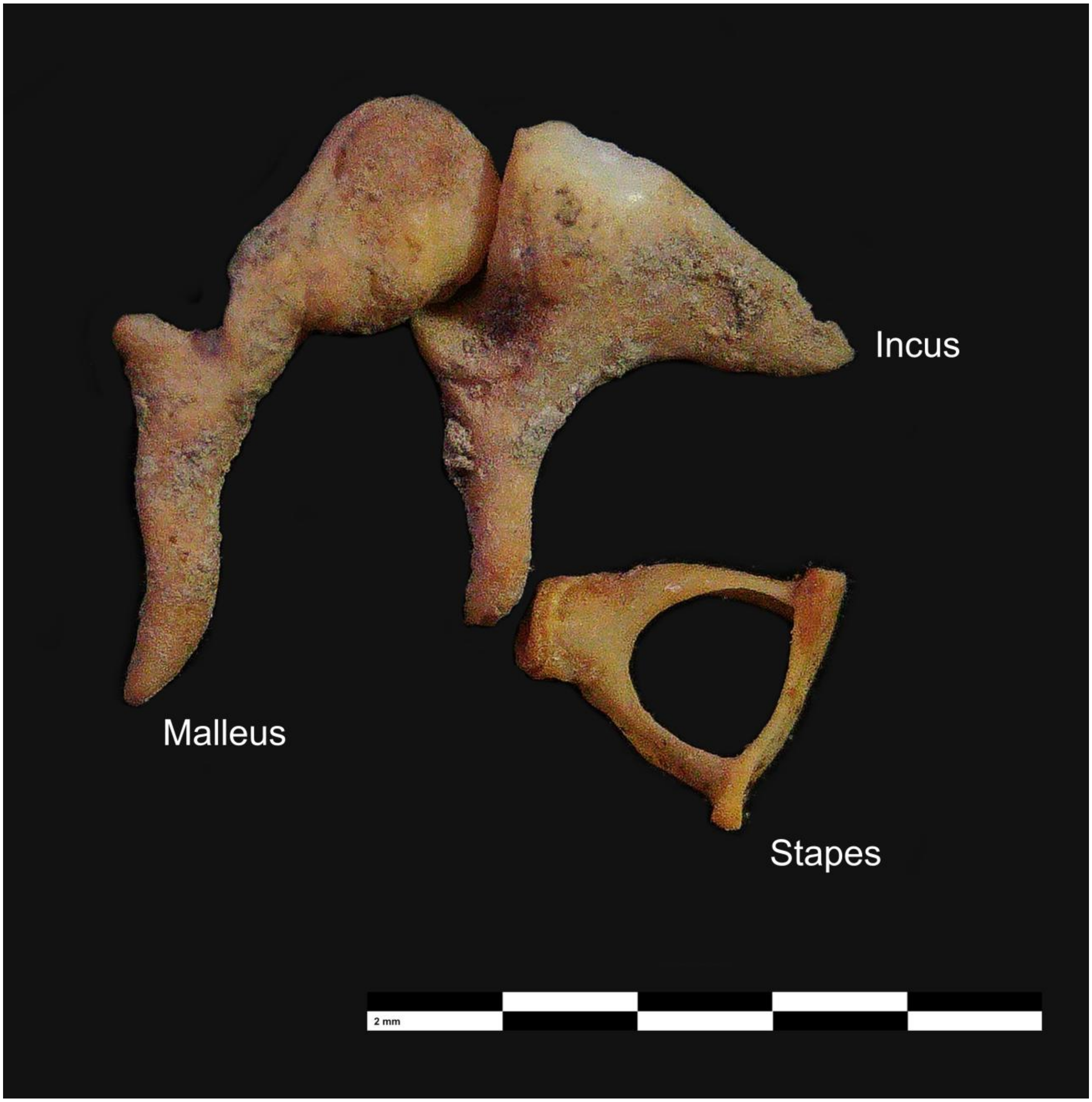


**Supplementary Figure 1:** Representative auditory ossicle bones used in this study. Each black and white segment in the scale bar represents 2 millimeters.

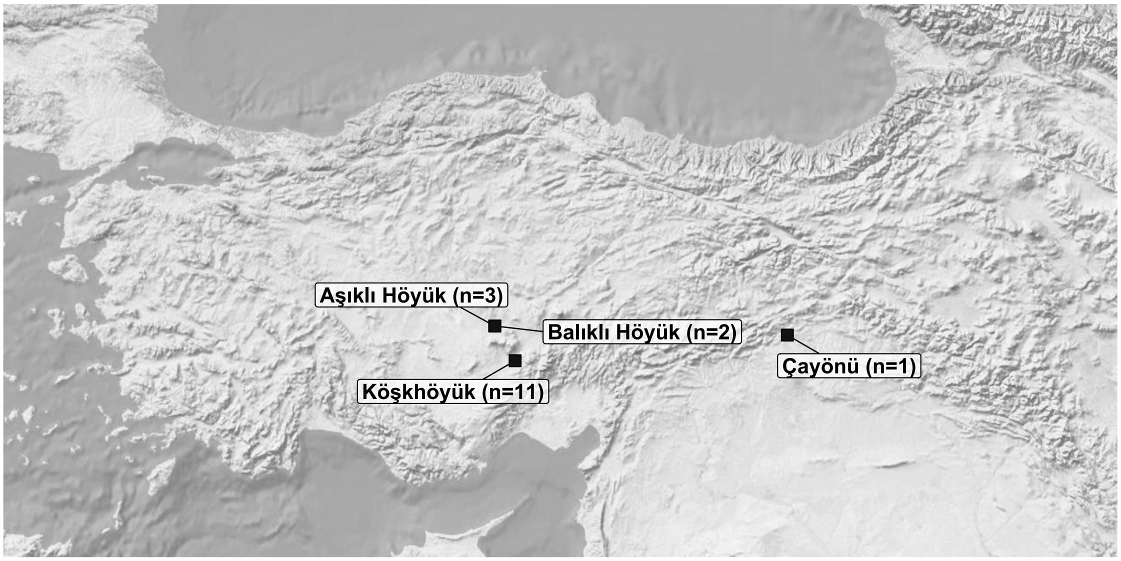


**Supplementary Figure 2:** Distribution of samples with both ossicle and petrous libraries. Sample size for each site is indicated in parentheses.

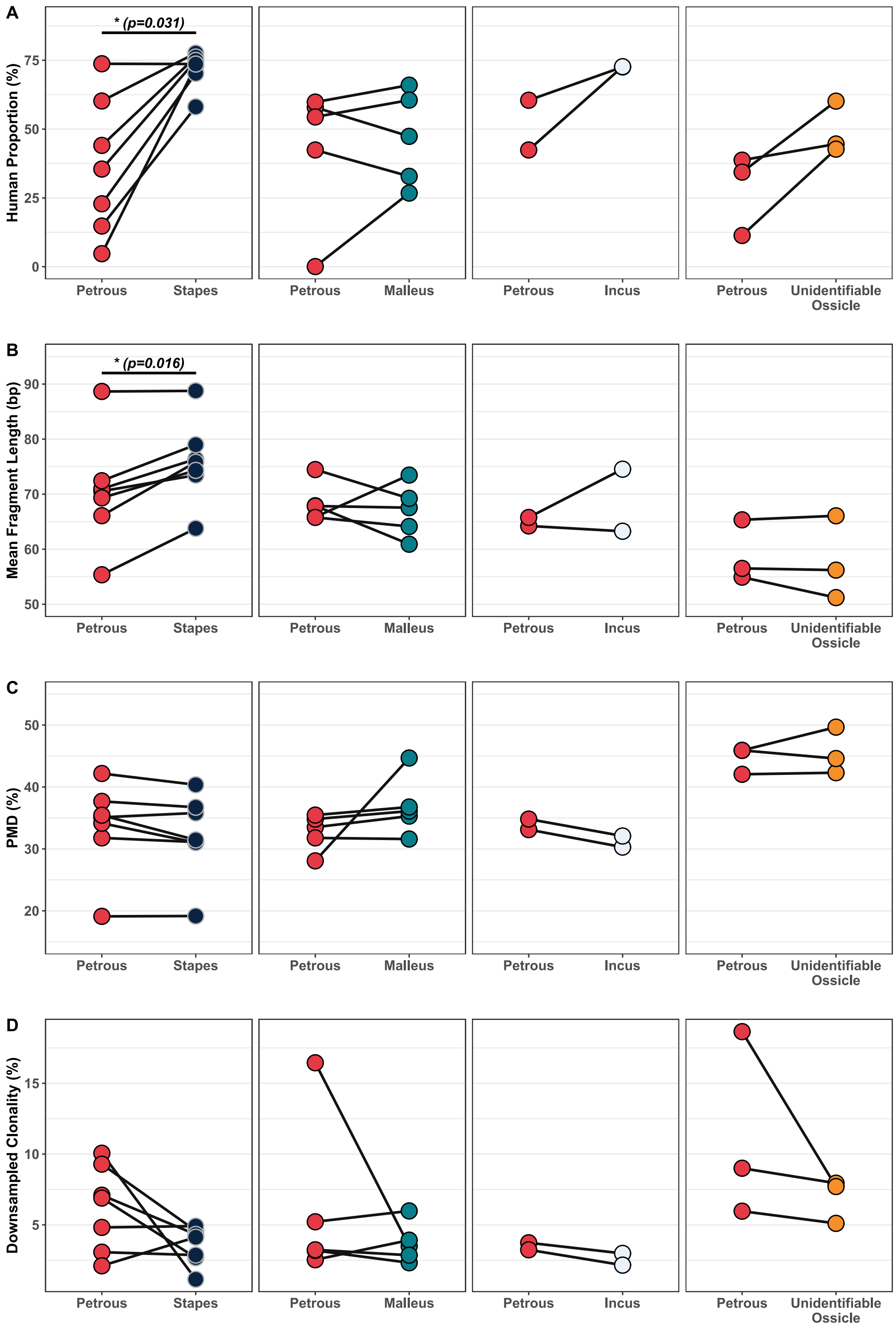


**Supplementary Figure 3:** Sequence statistics of ossicle and petrous libraries prepared from the same individuals. Wilcoxon signed-rank tests are indicated above the plots. Only statistically significant results are shown.
